# *Nubp2* is required for cranial neural crest survival in the mouse

**DOI:** 10.1101/656652

**Authors:** Andrew DiStasio, David Paulding, Praneet Chatuverdi, Rolf W. Stottmann

**Affiliations:** Divisions of Human Genetics, University of Cincinnati, Cincinnati, OH, 45229, USA; Developmental Biology, Cincinnati Children’s Hospital Medical Center, University of Cincinnati, Cincinnati, OH, 45229, USA; Department of Pediatrics, University of Cincinnati, Cincinnati, OH, 45229, USA

**Keywords:** Nubp2, neural crest, ENU mutagenesis, cilia

## Abstract

The N-ethyl-N-nitrosourea (ENU) forward genetic screen is a useful tool for the unbiased discovery of novel mechanisms regulating developmental processes. We recovered the *dorothy* mutation in such a screen designed to recover recessive mutations affecting craniofacial development in the mouse. *Dorothy* embryos die prenatally and exhibit many striking phenotypes commonly associated with ciliopathies, including a severe midfacial clefting phenotype. We used exome sequencing to discover a missense mutation in Nucleotide Binding Protein 2 (*Nubp2*) to be causative. This finding was confirmed with a complementation analysis between the *dorothy* allele and a *Nubp2* null allele (*Nubp2^Null^*). We demonstrate that *Nubp2* is indispensable for embryogenesis. NUBP2 is implicated in both the Cytosolic Iron/Sulfur cluster Assembly (CIA) pathway and in the negative regulation of ciliogenesis. Conditional ablation of *Nubp2* in the neural crest lineage with *Wnt1-cre* recapitulates the *dorothy* craniofacial phenotype. Using this model, we found that the proportion of ciliated cells in the craniofacial mesenchyme was unchanged, and that markers of the Shh, Fgf, and Bmp signaling pathways are unaltered. Finally, we show that the phenotype results from a marked increase in apoptosis within the craniofacial mesenchyme.

**Summary Statement:** An ENU screen identifies a novel allele of *Nubp2* which is then demonstrated to be required for cranial neural crest survival and proper midfacial development.

## Introduction

Vertebrate craniofacial development is a dynamic process highly dependent on a fascinating cell population known as the neural crest. Craniofacial neural crest cells (CNCC) are a multipotent and migratory cell type which delaminate from the dorsal aspect of the closing neural tube and travel ventrolaterally into the frontonasal process (FNP) and pharyngeal arches (PAs) where they become the mesenchyme of the emerging facial primordia [1, 2]. On each side of the mouse face, proliferation of the post-migratory neural crest causes medial and lateral nasal processes (MNP, LNP) to swell from the FNP around embryonic day (E) 10, resulting in paired horseshoe-shaped outgrowths with visible depressed nasal pits in the center. The outgrowth of these processes is coupled with that of the maxillary processes (MxPs) to bring the three primordia into contact [3, 4]. This contact and the subsequent fusion of the MNP, LNP, and MxP create the nostrils and upper lip [5]. As development continues past E10.5, further growth brings the paired MNPs into contact at the midface, where they fuse and give rise to the nasal septum, philtrum, and premaxilla.

During these early stages of craniofacial development, the CNCC rely on signaling cues from surrounding tissues to regulate their survival and the proliferation required to properly expand the facial primordia. Sonic Hedgehog (SHH) and Fibroblast Growth Factor 8 (FGF8) secreted by the surface ectoderm maintain CNCC survival and induce their proliferation [6–9]. Loss of these signals, or an inability of CNCC to properly transduce ligand binding within the cell, impairs the outgrowth and subsequent fusion of the facial primordia, often ultimately leading to midline defects of the face [10]. The CNCC are also a particularly vulnerable cell population as evidenced by their marked susceptibility to ribosome biogenesis insults, sensitivity to the teratogenic effects of alcohol and many examples of genetic mutations leading to elevated CNCC apoptosis [11–14] [15–17].

Despite significant advances in understanding the signaling pathways and transcriptional networks involved in craniofacial developmental disorders, their etiology remains unclear in many cases. Unbiased forward genetics approaches in model organisms allow discovery of previously unexamined roles for genes and pathways in embryogenesis, providing new insights and angles for investigation [18]. Here we report on a new allele from an N-ethyl-N-nitrosourea (ENU) screen in the mouse that is integral to craniofacial development. We found the causal mutation to be in the gene *nucleotide binding protein 2* (*Nubp2*). *Nubp2* is a highly conserved P-loop NTPase first described in early sequencing and mapping of cDNA isolated from E7.5 mouse embryos [19]. In *Saccharomyces cerevisiae* it was identified as *cytosolic Fe-S cluster deficient* (*Cfd1*) in a screen for genes able to convert Iron regulatory protein 1 into c-aconitase [20] [21]. These and subsequent studies in yeast and animal cell lines established that *Nubp2* is an integral component of the cytosolic iron-Sulphur (Fe-S) cluster assembly (CIA) pathway where it acts along with its homologous binding partner *Nubp1* as a scaffold for transfer of the Fe-S cofactor to non-mitochondrial apoproteins [22–24]. Later studies revealed that knockdown of *Nubp1* alone, or of both *Nubp1* and *Nubp2*, led to excessive centrosome duplication *in vitro* [25]. Further work indicated that silencing of *Nubp1* and *Nubp2* in cell culture led to supernumerary cilia formation [26]. An ENU screen isolated a mouse allele of *Nubp1* with interrupted lung budding and centriole over-duplication [27].

This implication of *Nubp2* in regulation of centriole duplication and ciliary dynamics, and the importance of the primary cilia to intercellular signaling, led us to hypothesize dysregulation of ciliogenesis would lead to interruptions in crucial CNCC signaling pathways, resulting in loss of mitogenic and/or survival signals and subsequent midfacial fusion defects. On the contrary, here we demonstrate that loss of *Nubp2* from the CNCC does not lead to ciliogenesis or signaling defects, but causes a rapid onset of apoptosis throughout the craniofacial mesenchyme following CNCC migration which underlies the distinctive craniofacial presentation of the *Dorothy* mutant. This represents the first example of a role for CIA pathway components in midfacial development.

## Materials and Methods

### Animal husbandry

All animals were housed and cared for in accordance with the Guide for the Care and Use of Laboratory Animals (National Institutes of Health), and all animal-associated activity was approved by the Institutional Animal Care and Use Committee of the Cincinnati Children’s Hospital Medical Center (Protocol # IACUC2016-0098). All animals were monitored daily by registered veterinary technicians and additional care provided where necessary. Euthanasia was performed via Isoflurane sedation followed by dislocation of the cervical vertebrae and secondary euthanasia. Mice were housed in ventilated cages provided with automated water supply and Purina 5010 Laboratory Rodent Chow *ad libitum*. During timed mating, noon of the day a copulation plug was detected was considered embryonic day (E) 0.5 (E0.5).

### ENU mutagenesis

ENU mutagenesis was performed as previously published [18]. Wildtype C57BL/6J (B6; Jackson Laboratory, Bar Harbor ME) 6-to 8-week-old males were given a fractionated dose of ENU (three weekly dosages of 90-100 mg/kg ENU, Sigma). Mutagenized males were bred with Tg(Etv1-EGFP)^B2192 GSat^ females (FVB/N TaC background) as part of a larger effort to identify potential mutants in cortical neuron patterning [28]. Following a three-generation backcross breeding scheme [18], G3 litters were dissected at late embryonic stages to detect gross developmental anomalies. The allele was propagated with test crosses onto the FVB background (to facilitate mapping if needed) until the causal allele was identified.

### Variant detection

Exome sequencing was carried out in the CCHMC Genetic Variation and Gene Discovery Core Facility (Cincinnati, OH, USA). An exome library was prepared from genomic DNA using the NimbleGen EZ Exome V2 kit (Madison, WI, USA). Sequencing of the resulting exome library was done with an Illumina Hi Seq 2000 (San Diego, CA, USA). The Broad Institute’s GATK algorithm was used for alignment and variant detection. Variants were then filtered as described in Table 1. Mutations were validated by Sanger sequencing.

**Table 1:**
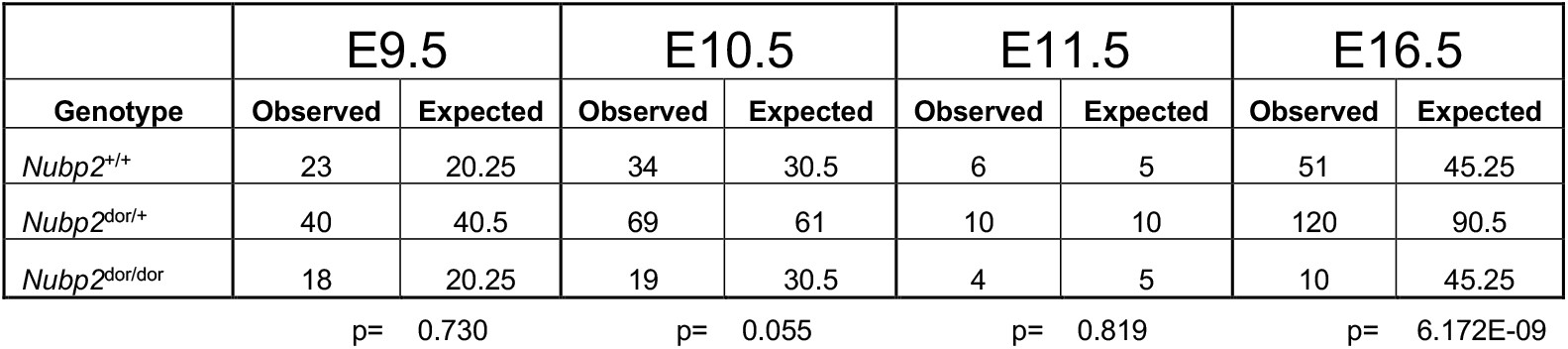
Recovery of *dorothy* mutants at different stages of embryogenesis

### Alleles and genotyping

*Nubp2*^Dor^ (*Nubp2* c.626T>A; Nubp2^V209D^; *dorothy*) was discovered in an ENU mutagenesis screen as described above and maintained on a mixed background. *Dorothy* genotyping was performed using the TaqMan™ Sample-to-SNP™ Kit using proprietary primers designed by the vendor targeting Nubp2:c:626T>A or Zbtb12:c:1061C>A (Thermo Fisher Scientific, Waltham, MA, USA). The *Nubp2*^tm1a^ (*Nubp2*^null^) allele was imported from the European Conditional Mouse Mutagenesis Program (EUCOMM, EM:08954) [29] using standard *in vitro* fertilization protocols. We excised the reporter gene trap cassette to create a traditional floxed allele *Nubp2*^tm1c^ (*Nubp2*^fx^) by crossing the *Nubp2*^tm1a^ with carriers of the Flippase enzyme [30]. PCR primers amplified a 470bp amplicon unique to the *Nubp2^tm1a^* allele (*Nubp2^tm1a^* F: TCATGCTGGAGTTCTTCGCC, *Nubp2^tm1a^* R: ATTCTTCCGCCTACTGCGAC). A separate primer pair amplify a 263bp sequence in the wildtype *Nubp2* allele and 457bp band in the *Nubp2*^fx^ allele (Nubp2WtFxF: GGATCCCAGTGCTGAGCTTT, Nubp2WtFxR: ACCACCTGCTAGCACTCAAC). For conditional ablations, we used the *Wnt1-cre* allele (B6.Cg-*H2afv*^*Tg(Wnt1-cre)11Rth*Tg(Wnt1-GAL4)11Rth^/J) [31] which was genotyped using a standard Cre detection PCR reaction (CreF: GCGGTCTGGCAGTAAAAACTATC, CreR: GTGAAACAGCATTGCTGTCACTT).

### Cell culture

Human Embryonic 293T (HEK293T) cells were cultured in Dulbecco’s modified eagle medium (DMEM, Sigma-Aldrich) supplemented with 10% Fetal Bovine Serum and 1% Penicillin-Streptomycin (Sigma-Aldrich). C- and N-terminal GFP-tagged *Nubp1* and *Nubp2* expression plasmids [32] were transfected using Lipofectamine^®^ 3000 (Thermo Fisher Scientific, Waltham, MA, USA) and imaged using a Nikon C2 Confocal microscope.

### Histology and Immunohistochemistry

Embryos designated for hematoxylin and eosin (H&E) staining were dissected and fixed overnight in 10% neutral buffered formalin solution (Sigma-Aldrich) at room temperature (RT) and then dehydrated in 70% ethanol for 48h. Following fixation and dehydration, samples were submitted to the CCHMC Pathology Research Core (Cincinnati, OH, USA) for paraffin embedding. Embedded samples were sectioned at a thickness of 10-14μM and processed through H&E staining. H&E slides were sealed with Cytoseal Mounting Medium (Thermo-Scientific). Samples designated for Immunohistochemical (IHC) analysis were dissected and fixed overnight in 4% paraformaldehyde at 4°C followed by immersion in 30% sucrose solution at 4°C for 24-48 hours until sufficiently dehydrated as observed by changes in sample buoyancy. Embryos were then embedded sagittally (E9.5) or coronally (E10.5) in OCT compound (Tissue-Tek) and cryosectioned at a thickness of 10-14μM. Sections were dried on a slide warmer at 42°C for 30-45 minutes for general IHC or dried at room temperature for 15-30 minutes when cilia were to be imaged. Samples were then soaked in boiling citrate buffer for 45-60 minutes for antigen retrieval followed by three PBS washes and blocked for 1 hour at RT in 4% normal goat serum in PBST. After blocking, sections were incubated overnight at 4°C with the following primary antibodies as indicated: CC3 (1:400, Cell Signaling 9661), PHH3 (1:500, Sigma H0412), AP2 (1:20, Developmental Studies Hybridoma Bank), Laminin (1:250, Sigma L9393), γ-Tubulin (1:1000, Sigma T6557), Arl13b (1:250, Proteintech 17711-1-AP). The following morning, sections were washed three times in PBS and incubated at RT in AlexaFluor 488 goat anti-mouse and AlexaFluor 594 goat anti-rabbit secondary (1:500, Life Technologies) for 1 hour at RT, washed 3 times in PBS and incubated in DAPI (1:1000, Sigma) for 20 minutes at RT. After 3 more PBS washes slides were sealed with ProLong Gold Antifade Reagent (Life Technologies) and imaged using a Nikon C2 confocal microscope. IHC images depicting cell death and proliferation were quantified using ImageJ (Rasband, W.S., ImageJ, U. S. National Institutes of Health, Bethesda, Maryland, USA, https://imagej.nih.gov/ij/, 1997-2018). Quantification of primary cilia was performed using NIS elements software (Nikon Instruments, Melville, NY, USA).

### In Situ Hybridization

Whole Mount In Situ Hybridization (WMISH) was performed as described previously [33] with a hybridization temperature of 65°C. Probes were transcribed from linearized plasmids using T7 RNA polymerase: *Etv5* (Xin Sun/ Gail Martin), *Ptc1* [34], *Msx1* [35], and *FGF8* [36].

### RNA Sequencing

3 WT and 3 *Wnt1-cre; Nubp2^cKO^* mutant E9.5 embryos were dissected from the yolk sac, and heads were snap frozen on dry ice. RNA was isolated and the individual samples were used for paired-end bulk-RNA sequencing (BGI-Americas, Cambridge, MA). RNA-Seq analysis pipeline steps were performed using CSBB [Computational Suite for Bioinformaticians and Biologists: https://github.com/csbbcompbio/CSBB-v3.0]. CSBB has multiple modules, RNA-Seq module is focused on carrying out analysis steps on sequencing data, which comprises of quality check, alignment, quantification and generating mapped read visualization files. Quality check of the sequencing reads was performed using FASTQC (http://www.bioinformatics.bbsrc.ac.uk/projects/fastqc). Reads were mapped (to mm10 version of Mouse genome) and quantified using RSEM-v1.3.0 [37].

## RESULTS

### Dorothy mutants were recovered in an ENU mutagenesis experiment

We performed a forward Ethyl-N-Nitrosourea (ENU) genetic screen in the mouse to discover and study unidentified alleles important for craniofacial development [18]. *Dorothy* (*dor*) mutants were isolated from this screen and readily identified by a characteristic craniofacial phenotype (Figure 1, Tables 1 and 2). At E16.5, *dorothy* mutants displayed a striking midfacial cleft phenotype in which the rostral-most nasal cartilage appears to be absent while the remaining nasal and maxillary structures were shifted laterally (Figure 1: A-F). We also noted micromelia and oligodactyly in each mutant embryo and commonly observed blood pooling in the distal hindlimbs reminiscent of the eponymous ruby slippers (Baum F. book) (Table 2). While all surviving embryos (6/6) collected at E16.5 had midline clefting and truncated limb phenotypes, mandibular development was variably affected (ranging from apparently normal development in 3/6 to complete agenesis in 1/6 non-necrotic E16.5 embryos observed). Curiously, despite the dysmorphic nature of the *dorothy* face, histological analysis revealed the presence of tooth buds in relatively appropriate positions, albeit with slightly delayed development (Figure 1: C, F, white dotted outlines). We also noted a significant portion of *dorothy* mutants did not survive beyond organogenesis stages. These commonly featured extensive hematomas and appeared morphologically to have arrested around E9.0-E9.5 (Figure 1: G-N, Table 1,2). During maintenance of the *dorothy* colony, living embryos past E10.5 appropriate for phenotypic analysis were increasingly rare. We hypothesize this was due to the increasingly inbred genetic background as the propagation of line was onto an FVB background in the event genetic mapping would be needed to clone the causal gene. The initial B6/FVB hybrid environment in the first matings may have led to increased *dorothy* mutant survival.

**Figure 1:**
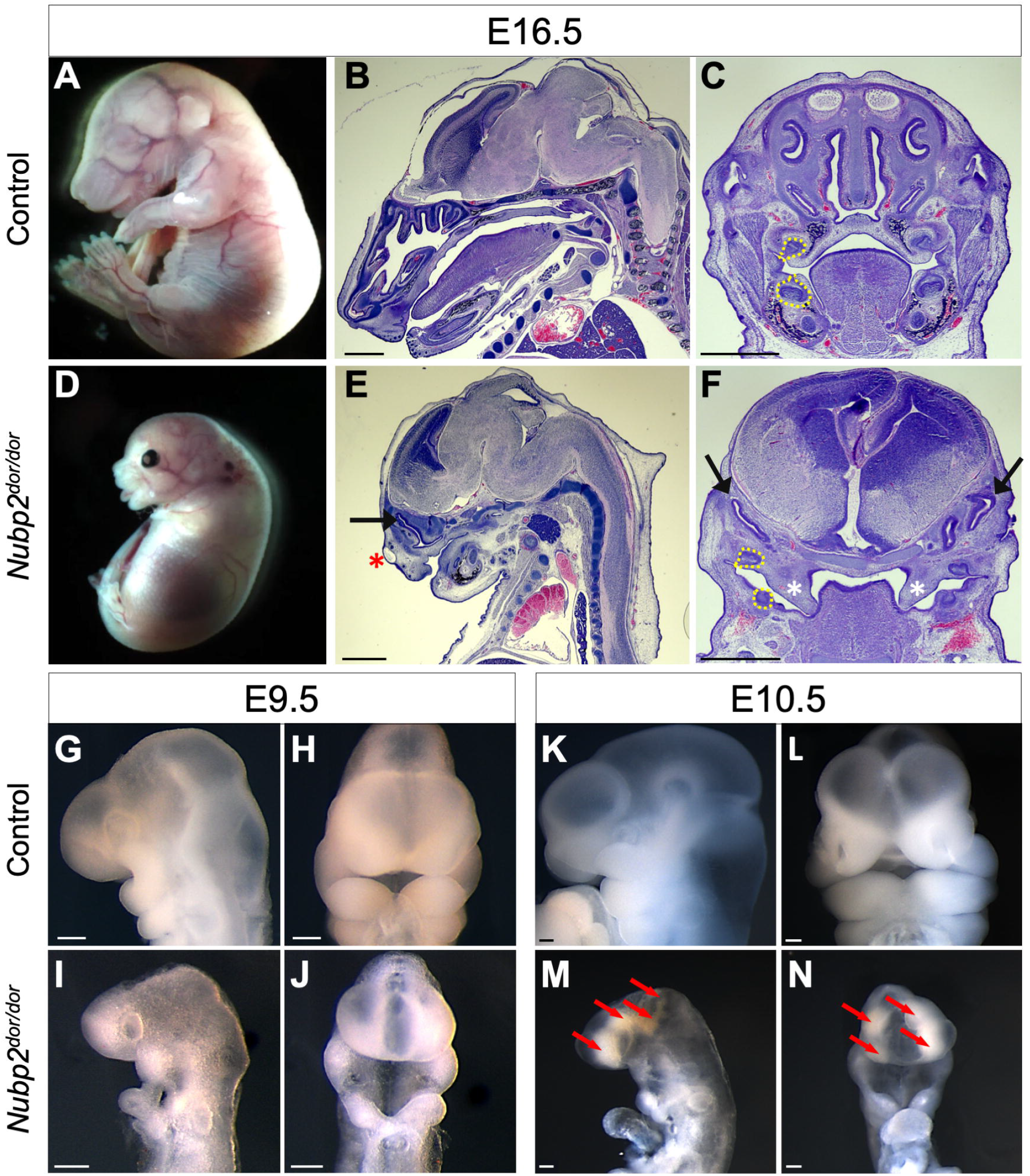
The Dorothy mutants have variable craniofacial phenotypes. Whole-mount and histological analysis of E16.5 control (A-C) and *dorothy* (D-F) mutant embryos. Mutants consistently present with midfacial clefting, micromelia and oligodactyly. The nasal cartilage (black arrows) is underdeveloped and ectopically located. Accumulation of blood or other fluid under the epithelium are also often noted (red asterisk). (F) Coronal sections also reveal cleft palate (black asterisks). Tooth buds are present but tooth development appears delayed (yellow dotted outlines). (G-N) *Dorothy* mutants with more severe phenotypes are developmentally arrested at the start of organogenesis (G-M), and often show signs of pervasive hematoma (red arrows). (Scale bars in B,C,E,F = 1mm, G-N = 200μm).

**Table 2:**
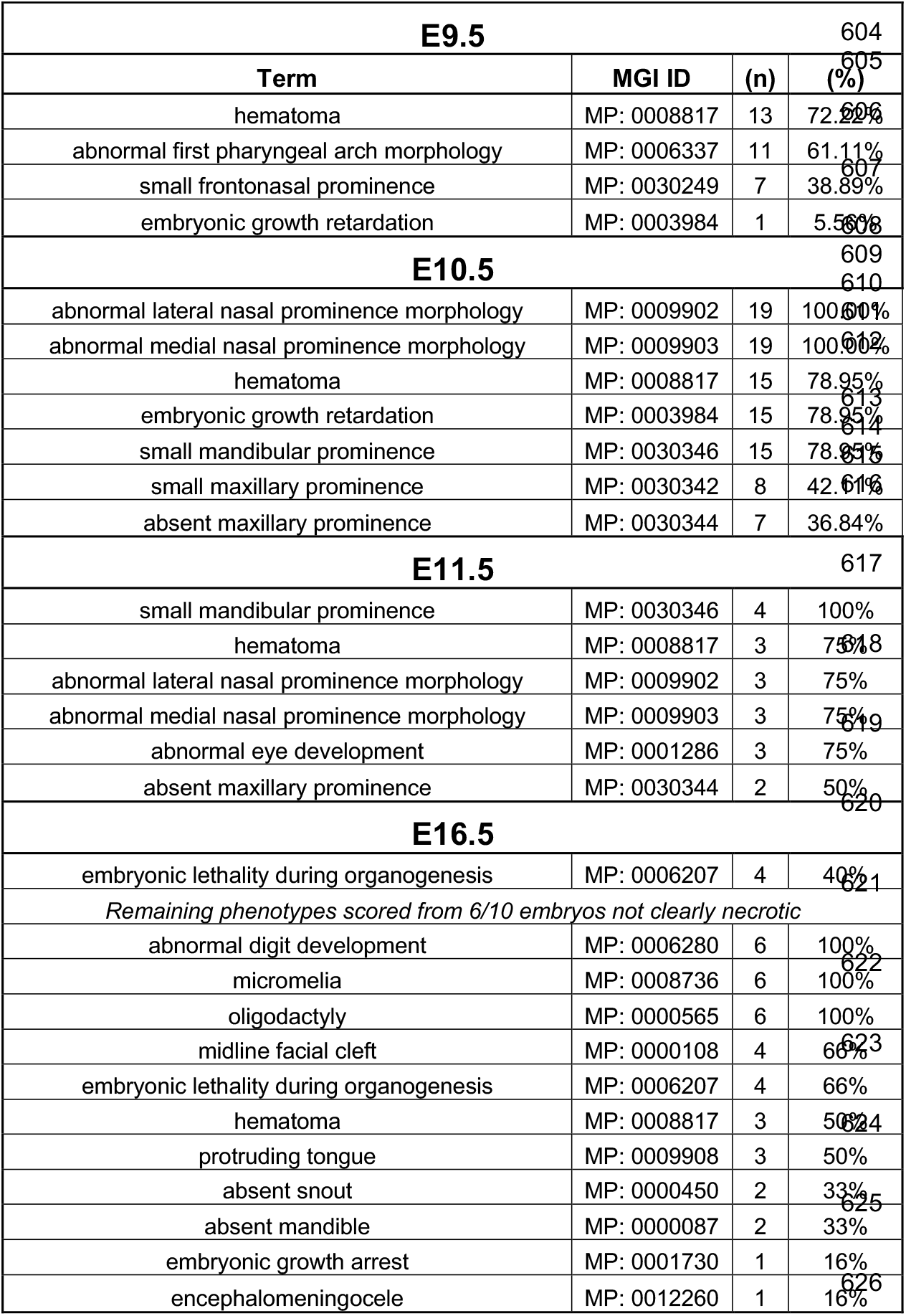
*Dorothy* phenotyping at various stages of embryogenesis

### The dorothy allele is a missense mutation in nucleotide binding protein 2 (Nupb2)

We performed exome sequencing to identify the causal allele in *dorothy* mutant. We sequenced three mutants and initially identified approximately 176,000 variants after quality control and read depth filters were applied. We first filtered out variants that were heterozygous in one or more mutants or that didn’t result from single nucleotide changes working under the assumption the *dorothy* allele is a recessive SNP, bringing the list of potential variants from ~176,000 to ~30,000 (Table 3). After filtering out SNPs known to be strain polymorphisms and variants predicted to have “low” impact, along with genes containing multiple variants (assumed to be highly polymorphic genes), 3 variants remained. One of these was in *Pbx2* which has been deleted in mouse with no resulting phenotype [38]. The remaining variants were in *nucleotide binding protein 2* (*Nubp2*) and *zinc finger and BTB domain containing 12* (*Zbtb12*). We further genotyped ten mutants and only homozygosity for the *Nubp2* was concordant with the phenotype (Table 3). The *Zbtb12* mutation was also homozygous in 9 unaffected embryos. Thus, Nubp2:c.626T>A; p.Val209Asp was identified as a candidate causal variant and this allele was designated *Nubp2^Dor^*.

**Table 3:**
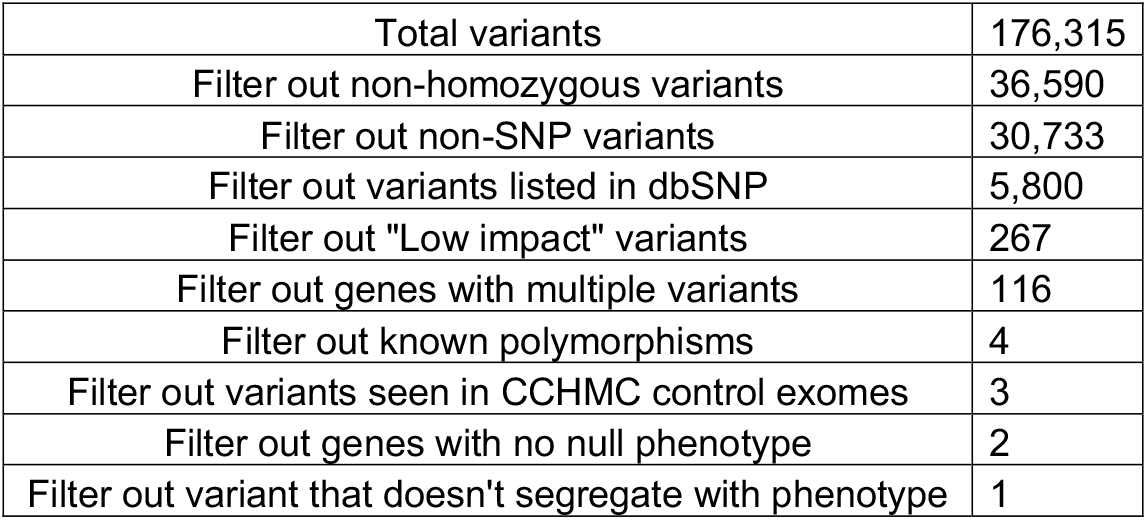
Explanation of exome sequencing result filtering

The *Nubp2^Dor^* missense mutation (Val209Asp, NUBP2^V209D^), changes the coding for a highly conserved nonpolar valine into a charged aspartic acid residue five amino acids downstream from a pair of Iron-interacting cysteine residues [23] (Figure 2A). Sanger sequencing confirmed the nucleotide change c626T>A in *dorothy* embryos (Figure 2B). Transfection of HEK293T cells with GFP-tagged h*Nubp2^wt^* and h*Nubp2^Dor^* expression constructs [32] demonstrated uninterrupted translation and cellular localization of hNUBP2^V209D^ (Figure 2C).

**Figure 2:**
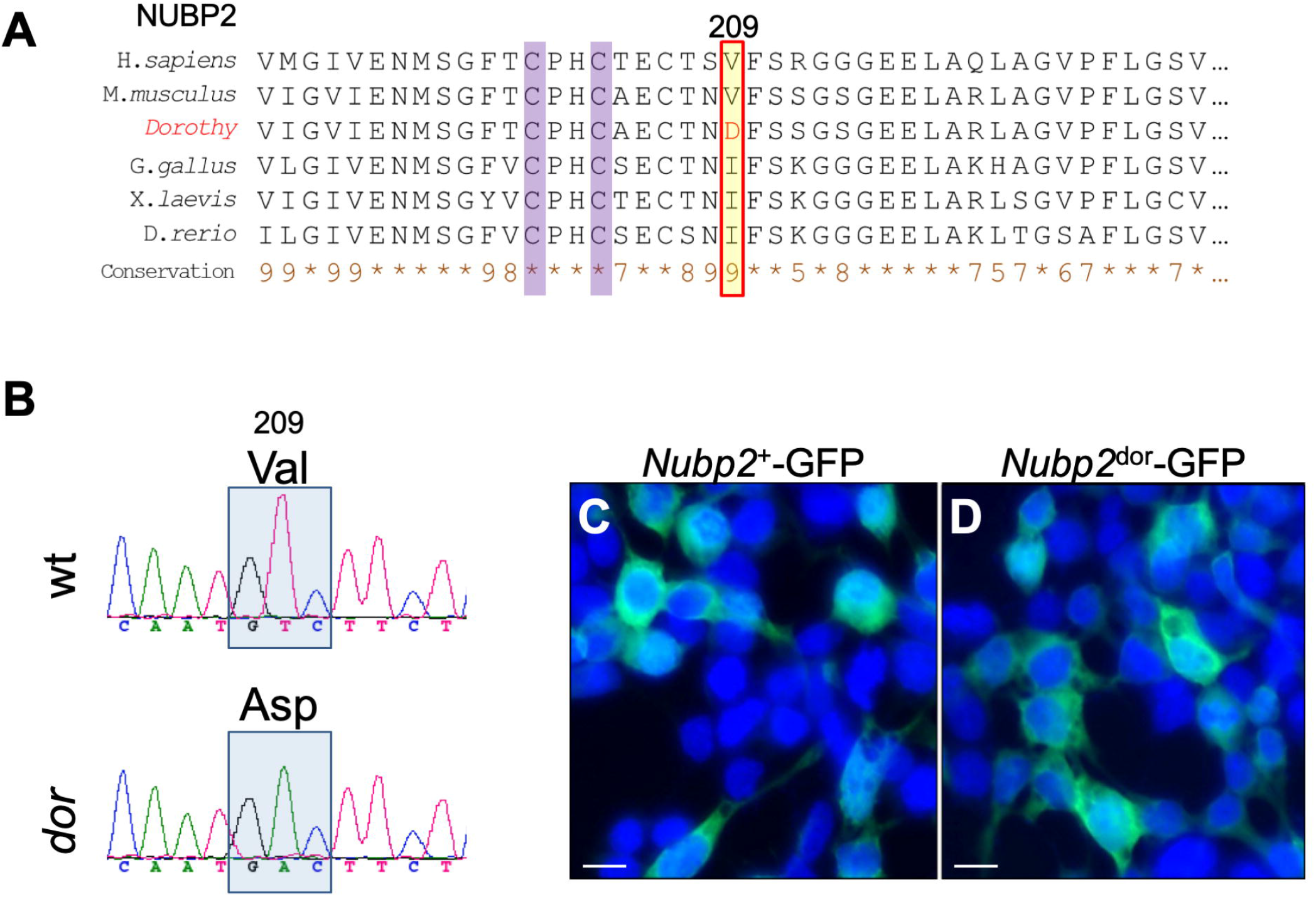
The *dorothy* mutation is an allele of *Nubp2*. (A) Amino acid sequence of NUPB2 in multiple model organisms. Nonpolar side chains (valine and isoleucine) at the position of the affected residue (red box, highlighted in yellow) are heavily conserved and located in close proximity to known functional Iron-interaction sites (highlighted in purple). In the *dorothy* mutant (p.Val209Asp; NUBP2^V209D^), this is replaced with an aspartate residue (red) harboring an electrically charged side chain. (B) Sanger sequencing confirms that Nubp2:c.626T>A is homozygous in embryos with the *dorothy* phenotype. (C,D) transfection of HEK293T cells with pAcGFP-C1-hNUBP2 (C) and pAcGFP-C1-hNUBP2^DOR^ (D) expression constructs revealed that the corresponding fusion proteins were both expressed and localized in roughly the same pattern, suggesting that the *Nubp2^dor^* mutation does not preclude translation of NUBP2 protein.

While the *dorothy* phenotype was completely concordant with homozygosity for the NUBP2^V209D^ allele, we sought to further confirm that the causal allele was in *Nubp2* via a complementation test with an independently derived null allele of *Nubp2*. We imported the *Nubp2^tm1a^* (*Nubp2^Null^*) allele from the international mouse phenotyping consortium [29, 39]. We first demonstrated that the *Nubp2^Null/Null^* embryos were not viable and were never recovered at early organogenesis stages (Table 4). Consistent with this, we noted frequent resorptions during these experiments, consistent with an early developmental lethality phenotype. We then intercrossed *Nubp2^dor/wt^* with *Nubp2^Null/wt^* and never recovered *Nubp2^Null/dor^* embryos at E16.5 (Table 5). This failure of the null allele to complement the ENU allele is the strongest possible genetic evidence that *dorothy* is an allele of *Nupb2*.

**Table 4:**
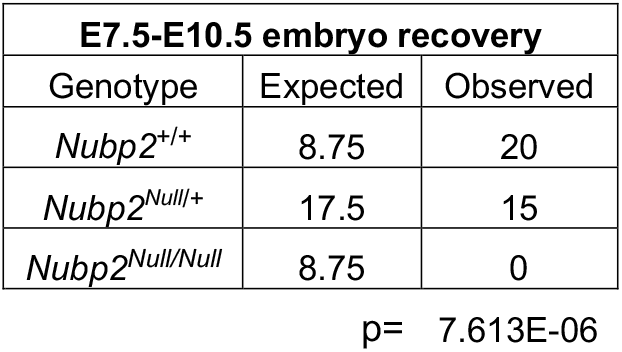
Embryonic lethality of *Nubp2* ablation

**Table 5:** Complementation test of *Nubp2^dor^* as the causative allele

| Embryo recovery |  |  |  |  |
| --- | --- | --- | --- | --- |
| Genotype | E16.5 |  | E10.5 |  |
|  | Expected | Observed | Expected | Observed |
| *Nubp2*<sup>+/+</sup> | 4.25 | 6 | 7.25 | 14 |
| *Nubp2*<sup>Null/+</sup> | 4.25 | 3 | 7.25 | 8 |
| *Nubp2*<sup>dor/+</sup> | 4.25 | 8 | 7.25 | 7 |
| *Nubp2*<sup>Null/dor</sup> | 4.25 | 0 | 7.25 | 0 |

p= 0.0344 p= 0.00347

### Nubp2 deletion from the neural crest lineage recapitulates the dorothy craniofacial phenotypes

Given the *dorothy* craniofacial phenotype, we hypothesized a unique requirement for *Nubp2* in the CNCC lineage. In order to test the hypothesis of cell-autonomous requirement for *Nubp2* in the CNCC and circumvent the early lethality and phenotypic variability of the *dorothy* allele, we used *Wnt1-cre* to ablate *Nubp2* in just the CNCC [31]. We first generated the *Nubp2^tm1c^* (*Nubp2^fx^*) allele by crossing the *Nubp2^tm1a^* with a FLP transgenic [30]. We then mated *Nubp2^Null/wt^* with *Wnt1-cre* to generate *Wnt1-cre Nubp2^Null/wt^*. These were again mated to *Nubp2^fx/wt^* or *Nubp2^fx/fx^* to create the *Wnt1-cre*; *Nubp2^cKO^*. We also perfomed similar crosses to generate *Wnt1-cre*; *Nubp^fx/fx^*. The *two* genotypes did not results in significantly different phenotypes and are collectively referred to here as *Wnt1-cre*; *Nupb2^cKO^* (Supplemental Table 1). The resulting *Wnt1-cre Nupb2^cKO^* embryos had phenotypes remarkably similar to the *dorothy* homozygous mutants (Figure 3A-H). At E15.5, embryos were recovered with midfacial clefts and undersized nasal, mandibular, and maxillary structures. Histological imaging revealed severe hypoplasia of the cartilage primordia of the nasal septum along with agenesis of the lateral cartilage primordia of the nasal capsule (Figure 3C,D,G,H, white and yellow dotted outlines, respectively). Combined with the cleft palate, this seems to lead to a situation where the olfactory epithelium is continuous with the oral cavity and, as in the *dorothy* mutant, the brain is shifted rostrally. We next set out to determine the developmental stage at which mutants are first phenotypically distinct from littermates. Virtually all *Wnt1-cre*; *Nubp2^cKO^* embryos (34/35) were anatomically indistinguishable from littermate controls at E9.5 (Figure 3: I-L; Table 6). In contrast, a day later at E10.5, a majority (14/17) of conditional mutants are easily identified by their conspicuously undersized facial primordia (Figure 3: M-P; Table 5). Thus, the *Wnt1-cre*; *Nubp2^cKO^* phenotype appears to emerge after CNCC migration, during early expansion stages, and results in a loss of craniofacial tissue.

**Figure 3:**
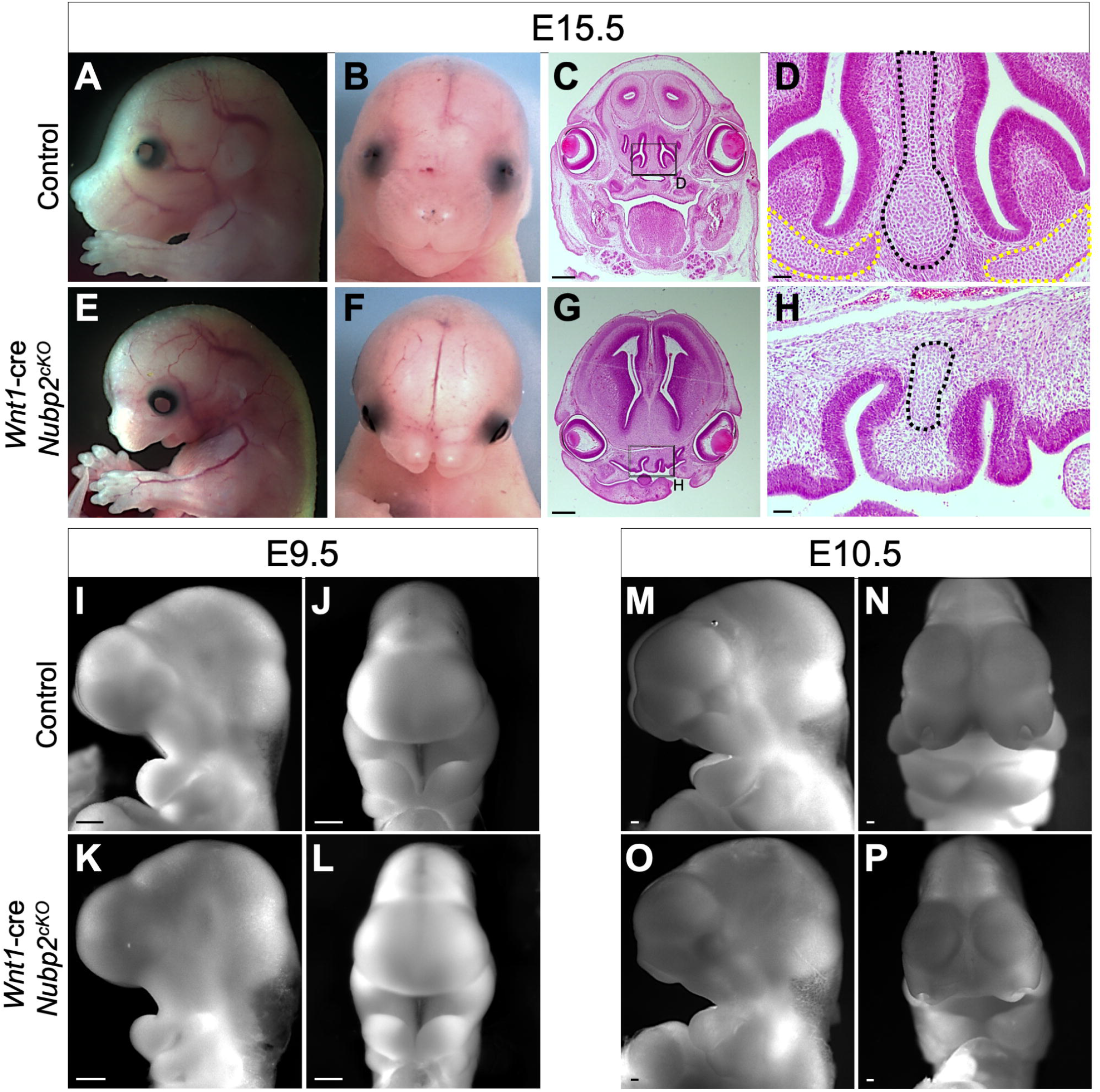
Conditional ablation of *Nubp2* in the neural crest lineage causes midfacial defects and recapitulates the craniofacial phenotype of the *dorothy* mutant. (A-H) *Wnt1-cre*; *Nubp2^cKO^* embryos recovered at E15.5 have severe craniofacial abnormalities including midline clefting and mandibular hypoplasia. (C, G) H&E stained coronal sections demonstrate palatal shelves fail to elevate. (D, H) higher magnification reveals that the cartilage primordium of the nasal septum is severely truncated (black dotted outline), while cartilage primordium of the nasal capsule lateral to the septum (yellow dotted outline) appears to be completely absent in the mutant. (I-L) At E9.5 *Wnt1-cre*; *Nubp2^cKO^* mutants are indistinguishable from wild-type (M-P) but at E10.5 the phenotype becomes apparent with hypoplastic nasal prominences, maxillary and mandibular prominences). (Scale bars in C, G = 500μm; D,H=50μm; I-P 100μm.)

**Table 6:** Phenotypic description of *Wnt1-cre*; *Nubp2*^cKO^ embryos

| E9.5 |  |  |  |
| --- | --- | --- | --- |
| Term | MGI ID | (n) | (%) |
| no abnormal phenotype detected | MP: 0002169 | 34/35 | 97% |
| abnormal first pharyngeal arch morphology | MP: 0006337 | 1/35 | 3% |
| E10.5 |  |  |  |
| abnormal lateral nasal prominence morphology | MP: 0009902 | 14/17 | 82.35% |
| abnormal medial nasal prominence morphology | MP: 0009903 | 14/17 | 82.35% |
| small maxillary prominence | MP: 0030342 | 12/17 | 70.59% |
| small mandibular prominence | MP: 0030346 | 4/17 | 28.57% |
| no abnormal phenotype detected | MP: 0002169 | 3/17 | 17.65% |
| embryonic growth retardation | MP: 0003984 | 1/17 | 5.88% |
| E11.5 |  |  |  |
| abnormal lateral nasal prominence morphology | MP: 0009902 | 7/9 | 77.78% |
| abnormal medial nasal prominence morphology | MP: 0009903 | 7/9 | 77.78% |
| small maxillary prominence | MP: 0030342 | 7/9 | 77.78% |
| absent mandible | MP: 0000087 | 6/9 | 66.67% |
| hematoma | MP: 0008817 | 4/9 | 44.44% |
| no abnormal phenotype detected | MP: 0002169 | 2/9 | 22.22% |

### Deletion of Nubp2 in the CNCC does not affect centriole duplication or ciliogenesis

Due to previously described roles for *Nubp2* in regulation of centriole duplication and ciliogenesis [25, 26], we hypothesized that similar disruptions in primary cilia biology may compromise the CNCC-derived mesenchyme. We used immunohistochemistry with confocal fluorescence microscopy to image and quantify centrosomes and primary cilia within the E9.5 craniofacial mesenchyme (Figure 4). We used γ-tubulin to mark centrioles and Arl13b to highlight the ciliary axoneme. The cilia in *Wnt1-cre*; *Nubp2^cKO^* mutants were qualitatively indistinct from those in littermates (Figure 4, A-H). We quantified the number of cilia and centrioles per nucleus for conditional mutants and controls. We observed slight increase in average cilia/cell (Figure 4I, p=0.009) but no difference in centriole number (Figure 4J; p=0.422).

**Figure 4:**
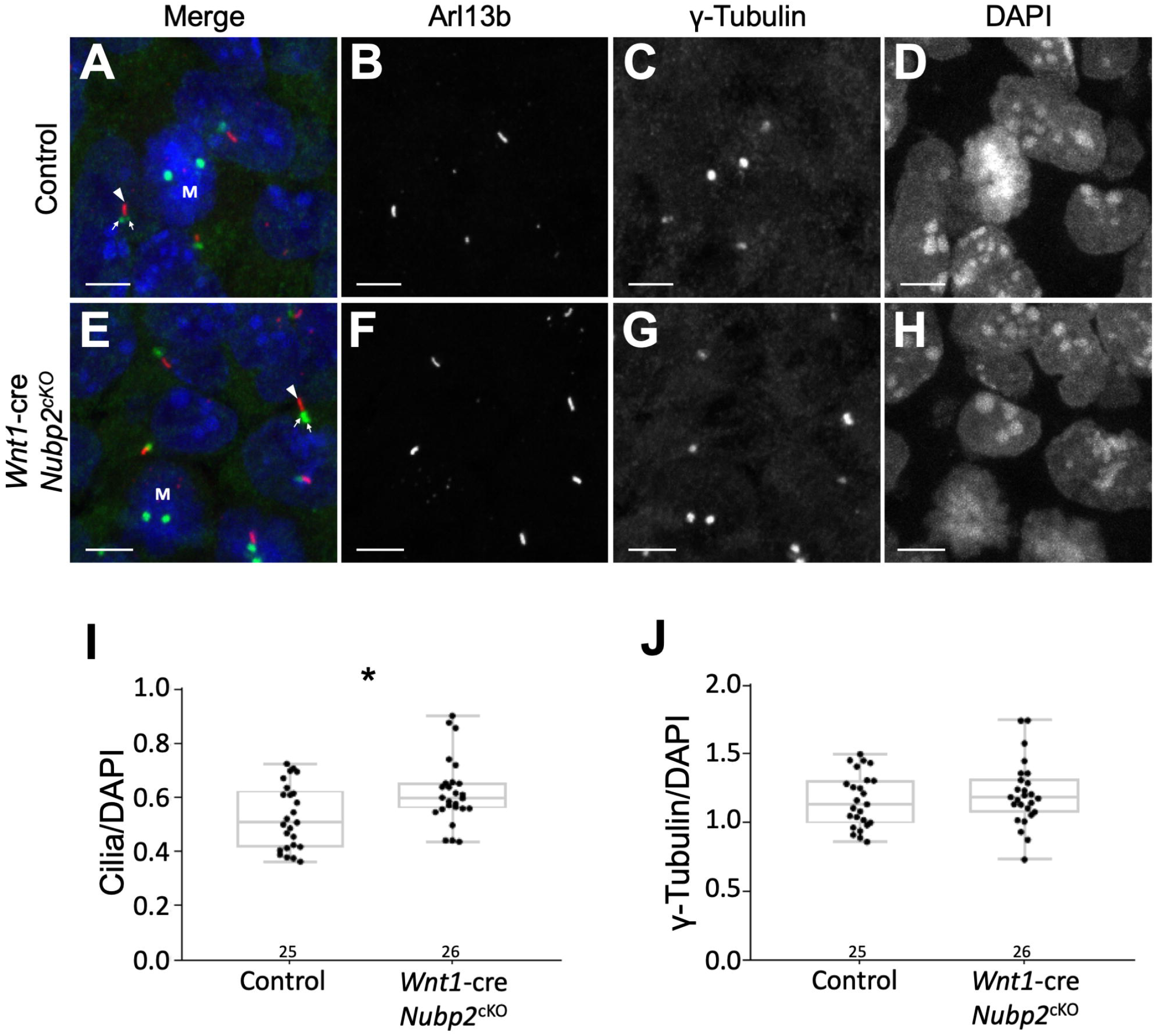
Craniofacial mesenchymal cells have normal primary cilia and centrioles. (A-H) representitive fields showing immunostaining with fluorescent antibodies against γ-Tubulin and Arl13b in E9.5 craniofacial mesenchyme. No gross abnormalities are observed in mutant cilia (white arrows) or centrioles (white arrowheads). Mitotic cells can be observed with two centrioles (white M). (I) a minor, but statistically significant increase in the proportion of cilia per cell was measured in mutants. (J) There was no observed difference in the proportion of centrioles per cell in mutants. Scale bars: 5μm.

### Loss of Nupb2 does not alter Shh, Fgf or Bmp signaling in craniofacial neural crest cells

The midfacial defects found in the *Wnt1-cre*; *Nubp2^cKO^* mutants suggested a number of developmental signaling pathways may be adversely affected. Appropriate levels of Shh, Fgf and Bmp signaling are all known to be crucial for proper CNCC development. We addressed each of these individually with whole mount in situ hybridization on selected markers as canonical transcriptional targets of each signaling pathway.

Hedgehog signaling is well known to be a survival signal for the CNCC [40]. Genetic ablation of smoothened (*Smo*) in the neural crest using the same *Wnt1-cre* allele used here led to a phenotype highly reminiscent of the *dorothy* phenotype we present here [6]. However, *Ptch1* expression was qualitatively and unaffected in *Wnt1-cre*; *Nubp2^cKO^* mutants at E9.5 (Figure 5, A-D). Fgf signaling is required from the surface ectoderm to the neural crest-derived facial mesenchyme during early craniofacial development [8, 9]. *Etv5* as a measure of Fgf signaling was transcribed normally (Figure 5, E-H). Finally, disturbances in bone morphogenetic protein (Bmp) signaling during craniofacial development have been associated with defective fusion of the facial primordia [41], and mutation of the downstream transcription factor *Msx1* causes cleft lip/palate. *Msx1* levels also appeared unchanged in conditional mutants (Figure 5, I-L).

**Figure 5:**
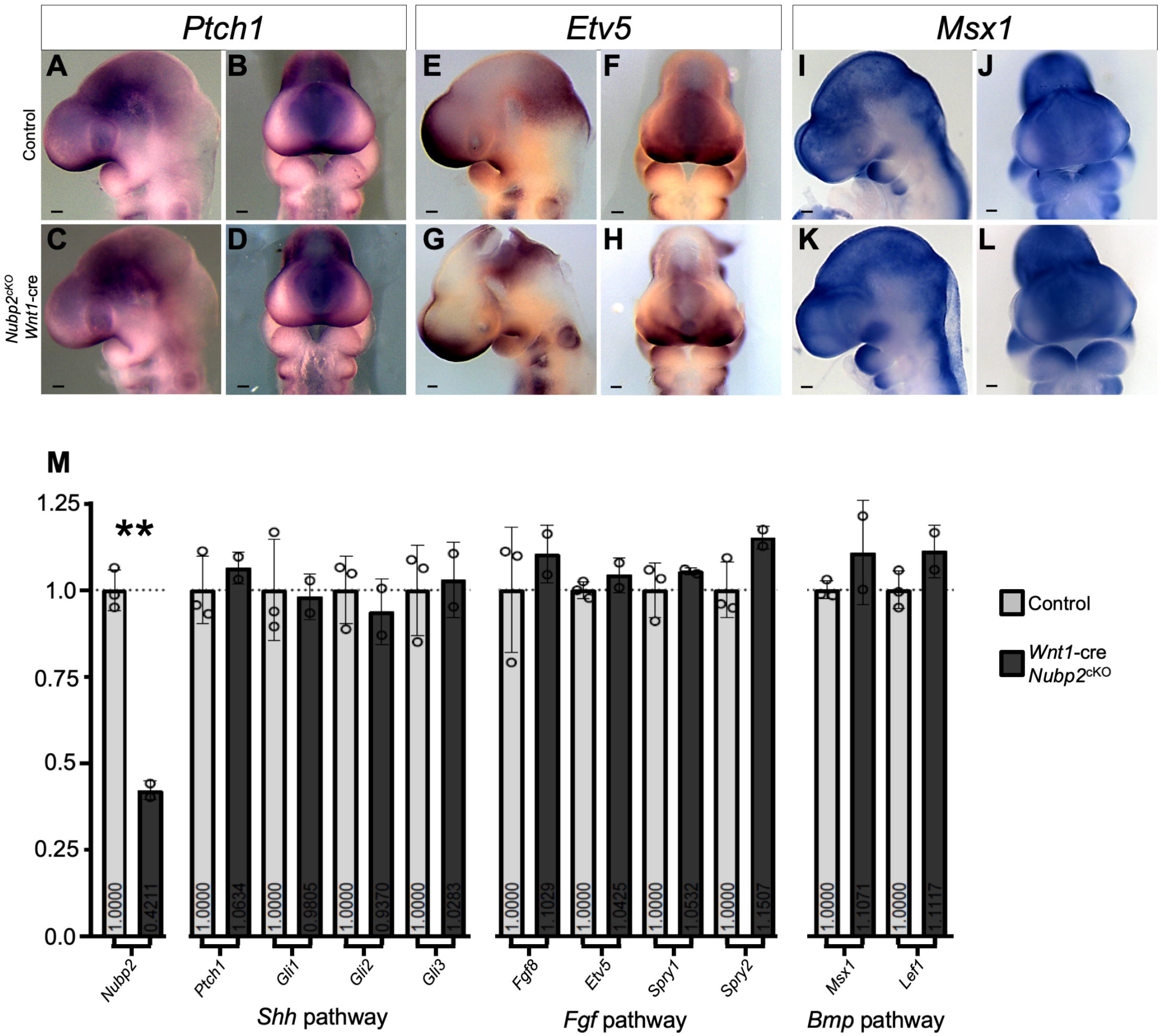
Major developmental signaling pathways are unaffected in *Wnt1-cre*; *Nubp2^cKO^* embryos. (A-L) Whole mount in situ hybridizations of E9.5 embryos and (M) normalized FPKM values derived from RNA-sequencing of dissected E9.5 heads show that the Shh (A-D), Fgf (E-H) and Bmp (I-L) signaling pathways are unaltered at E9.5 in *Wnt1-cre*; *Nubp2^cKO^* embryos (Scale bars =100μm).

In order to look for more markers of these canonical pathways and perform an unbiased transcriptional analysis of mutants, we dissected E9.5 heads from three controls and two mutants and performed bulk RNAsequencing. We first measured levels of Shh signaling (*Ptch1, Gli1, Gli2, Gli3*), FGF signaling (*Fgf8, Etv5, Spry1, Spry2*), and Bmp signaling (*Msx1, Lef1*) and saw no significant changes between wild-type and mutant embryos (Figure 5 M).

### Neural crest-specific ablation of Nubp2 does not significantly alter proliferation, but increases apoptosis

In the absence of intercellular signaling defects, we hypothesized that the failure of facial primordia to sufficiently expand may be due to altered proliferation and/or cell survival. The craniofacial mesenchyme at this stage must be highly proliferative in order to produce the maxillary, mandibular, and nasal prominences and to ensure that the paired MNPs meet at the midline to form the intermaxillary segment. At E9.5 the ratio of proliferating cells in the craniofacial mesenchyme as measured by a mitotic index of Phosphorylated Histone 3 (PHH3) to DAPI-stained nuclei is unchanged in *Wnt1-cre*; *Nubp2^cKO^* mutants (Figure 6, A-C; p=0.453). This ratio is mildly elevated at E10.5, when the LNP and MNP are noticeably truncated in comparison to littermate controls (Figure 6, D-F; p=0.060).

**Figure 6:**
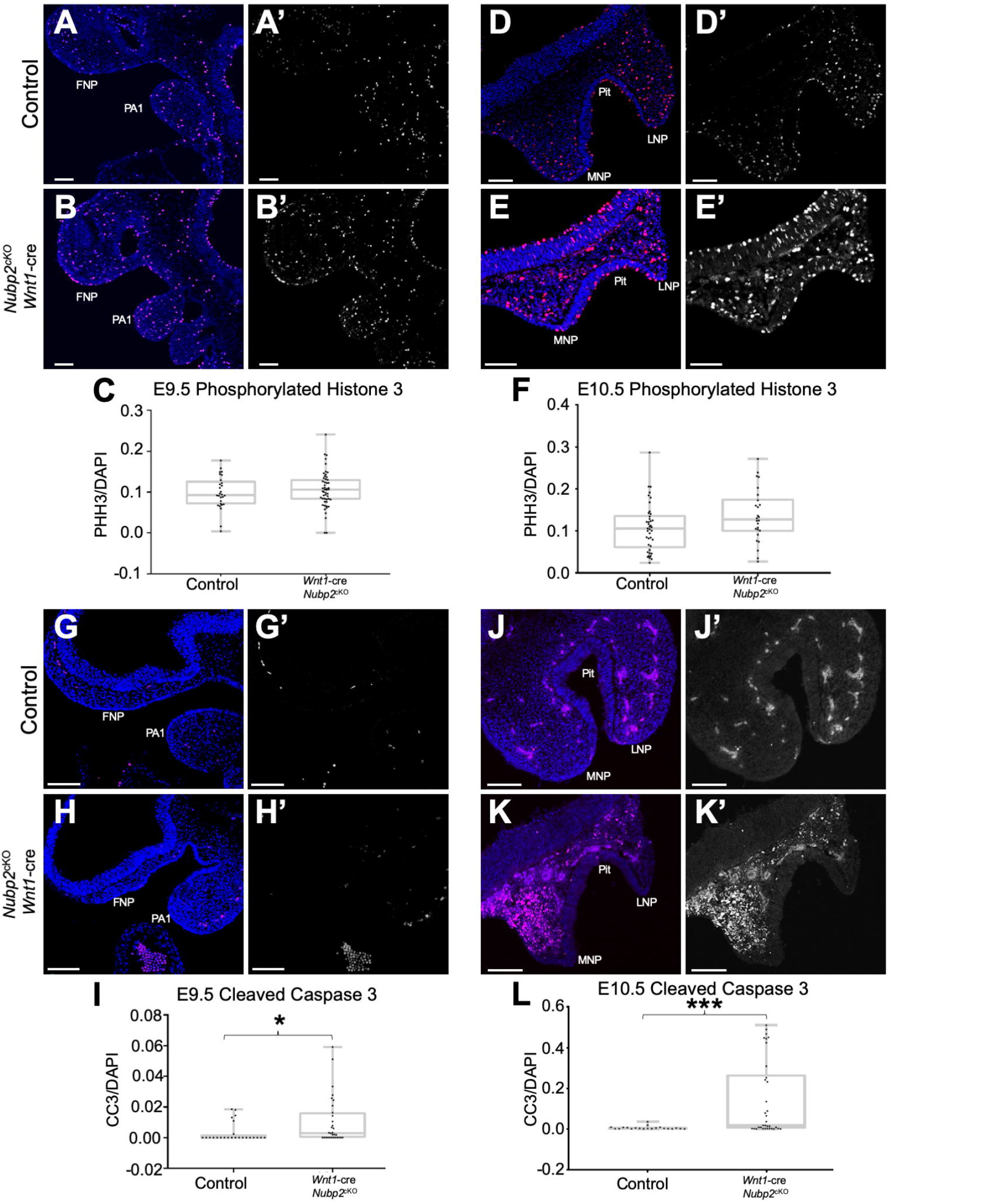
Loss of *Nubp2* leads to decreased survival of craniofacial neural crest. (A-E’) immunofluorescent staining for Phosphorylated Histone 3 (PHH3) at E9.5 (A-C; p=0.453) and E10.5 (D-F; p=0.060) shows no significant change in the proportion of proliferating *Wnt1-cre*; *Nubp2^cKO^* craniofacial mesenchymal cells at either stage. (G-L) in contrast, staining for cleaved caspase 3 (CC3) at E9.5 (G-I; p=0.040) shows a slight increase in apoptotic mesenchyme and by E10.5 (J-L; p<0.0001) there is a striking and statistically significant increase. Scale bars: 100μm.

We then assayed levels of apoptosis by calculating the ratio of Cleaved Caspase 3 (CC3) to DAPI-stained nuclei. We observed increased apoptosis at E9.5 (Figure 6, G-I; p=0.040) and a drastic increase at E10.5 (Figure 6, J-L; p<0.0001). This increase was more substantial in the MNPs than the LNPs (Fig. 6K) Taken together, these data show CNCC lacking *Nubp2* are capable of migrating into the FNP and PA’s, correctly regulating ciliogenesis, responding properly to intercellular signaling, and proliferating at a normal rate, but undergo abrupt apoptosis between E9.5 and E10.5.

## DISCUSSION

In this report, we describe our identification of the *dorothy* mutation and determine it to be an allele of *Nubp2*. The *dorothy* mutant has a variable phenotype with highly penetrant micromelia, syndactyly, dysmorphic nasal cartilage, and midfacial clefting (Figure 1, Tables 1, 2). Exome sequencing identified a variant in *Nubp2* as a candidate causal mutation (Figure 3, Table 3). We used a null allele to show *Nubp2* is required for early development (Table 4) and also used this allele in a complementation test to further demonstrate *dorothy* is a hypo-morphic allele of *Nubp2* (Table 5). We then determined that the *dorothy* craniofacial phenotype is due to a cell-autonomous requirement for *Nubp2* in the NCC lineage (*Wnt1-cre*; *Nubp2^cKO^*), with a phenotype emerging between E9.5 and E10.5 (Figure 3). We used these *Wnt1-cre*; *Nupb2^cKO^* embryos to show that ciliogenesis is only mildly affected in the craniofacial mesenchyme (Figure 4), despite previously demonstrated roles for *Nubp2* as a regulator of ciliogenesis [26]. Several canonical signaling pathways required for craniofacial development and the proportion of proliferating cells are unaltered in *Wnt1-cre*; *Nupb2^cKO^* embryos (Figure 5, 6). Ultimately, we observed a conspicuous increase in apoptosis among the post-migratory CNCC mesenchyme of the craniofacial primordia starting around E9.5 (Figure 6, G-L). We hypothesize that it is this ectopic cell death which prevents the craniofacial primordia from growing sufficiently to bring the opposing MNPs into contact, resulting in severe midfacial clefting and hypoplasia of the mandible, nasal septum and nasal capsule. This study is the first to demonstrate a requirement for *Nubp2* in development and shows a more specific role in craniofacial development.

The discovery of this role for *Nubp2* illustrates the continuing power of unbiased forward genetic screens, as it is unlikely the gene would have otherwise been selected for study in craniofacial development. To our knowledge, no clinical disorders have been associated with *Nubp2* variants. During the course of this study, the IMPC characterized the *Nubp2^tm1b/tm1b^* mouse as preweaning lethal with complete penetrance (*Nubp2^tm1b^* is derived from the *Nubp2^null^* allele we described above) [39]. ENU mutagenesis creates missense mutations that are often hypo-morphic, providing more insight into how essential genes function in specific developmental processes. For instance, ENU-induced mutations of *Hsd17b7* and *Grhl2* provided informative craniofacial and neural phenotypes despite the fact that complete deletion of either gene is lethal before completion of organogenesis [42–45]. In many such cases the effects of point mutations cannot be inferred from knockout studies alone, and forward screens can provide a genetic model more relevant to the natural population. For example, patients with mutations in both alleles of *POLR1C* present with Treacher Collins Syndrome, while *Plr1c^null/null^* mice are embryonic lethal [39, 46, 47].

Along with NUBP1, NUBP2 participates in an early step of the CIA pathway as a scaffold for the Fe/S cluster, which is transferred to target proteins via ATP hydrolysis [22, 23]. The NUBP1^NISW^ mutant is compromised in its ability to bind NUBP2, thus abrogating the canonical heterodimer scaffold [27]. This loss may explain similarities between the *Nubp1^Nisw^* and *dorothy* phenotypes, which include micromelia, syndactyly, and defects in ocular development. Cataracts are also associated with mutations in *Xpd/Ercc2* and *Spry2*, both of which are extramitochondrial Fe/S proteins [48–51]. Loss of *Iop1* in the mouse, which operates downstream of NUBP1/NUBP2 in the pathway, causes lethality prior to E10.5 as does loss of *Nubp2* [52, 53]. These data taken together suggest that aspects of the *dorothy* and *Nubp1^Nisw^* phenotypes may be attributed to decreased CIA pathway activity, but also that complete loss of the pathway may be lethal prior to organogenesis. This is consistent with our inability to recover *Nubp2^Null/Null^* embryos. It is possible that the NUBP2 homodimer is much more important for Fe/S cluster maturation in the craniofacial mesenchyme, or that the ATPase has acquired a secondary role during mammalian evolution.

Cases of “protein moonlighting” can be established when proteins acquire mutations or new post-translational modifications during evolution and new interactions arise [54]. One relevant example is the case of *Irp1*. This enzyme catalyzes the conversion of citrate to isocitrate when the CIA pathway provides an Fe/S cluster, but in the absence of cofactor IRP1, becomes an active RNA-binding protein modifying the turnover of target mRNAs [55]. Presumably these functions evolved as a way for the cell to couple the activity of an important Fe/S protein to regulation of transcripts relevant to iron homeostasis. Could *Nubp2* have another function outside of the CIA pathway that explains the presence of midfacial clefting in *Nubp2*, but not *Nubp1* mutants? Using the Harmonizome database [56] we found that both genes share 12 interaction partners, mostly other members of the CIA pathway such as CIAO1 and MMS19 [57–59]. Besides these, NUBP1 has 16, and NUBP2 43, unique interaction partners. Among the unique partners of each protein are several proteins involved in the regulation of apoptosis. Notably, among the unique partners of NUBP2 is Pleckstrin Homology-Like Domain Family A Member 3 (PHLDA3), a pro-apoptotic repressor of the oncogene *Akt* [60]. Our RNAseq experiment showed that *Phlda3* transcripts were significantly upregulated in E9.5 *Wnt1-cre*; *Nubp2^cKO^* heads, suggesting a potential functional significance for this interaction. NUBP2 (but not NUBP1) also has an interaction with E2F transcription factor 1 (E2F1), which can induce both proliferation and apoptosis [61]. The ChIP Enrichment Analysis (ChEA) database shows that E2F1 binds the promoter regions of both NUBP1 and NUBP2 [62]. In light of the increased apoptosis in the craniofacial mesenchyme of *Nubp2^cKO^* and in the developing lungs of *Nubp1^Nisw^* embryos, future studies could explore a role for the proteins in the regulation of cell death.

---

**Supplemental Table 1:**
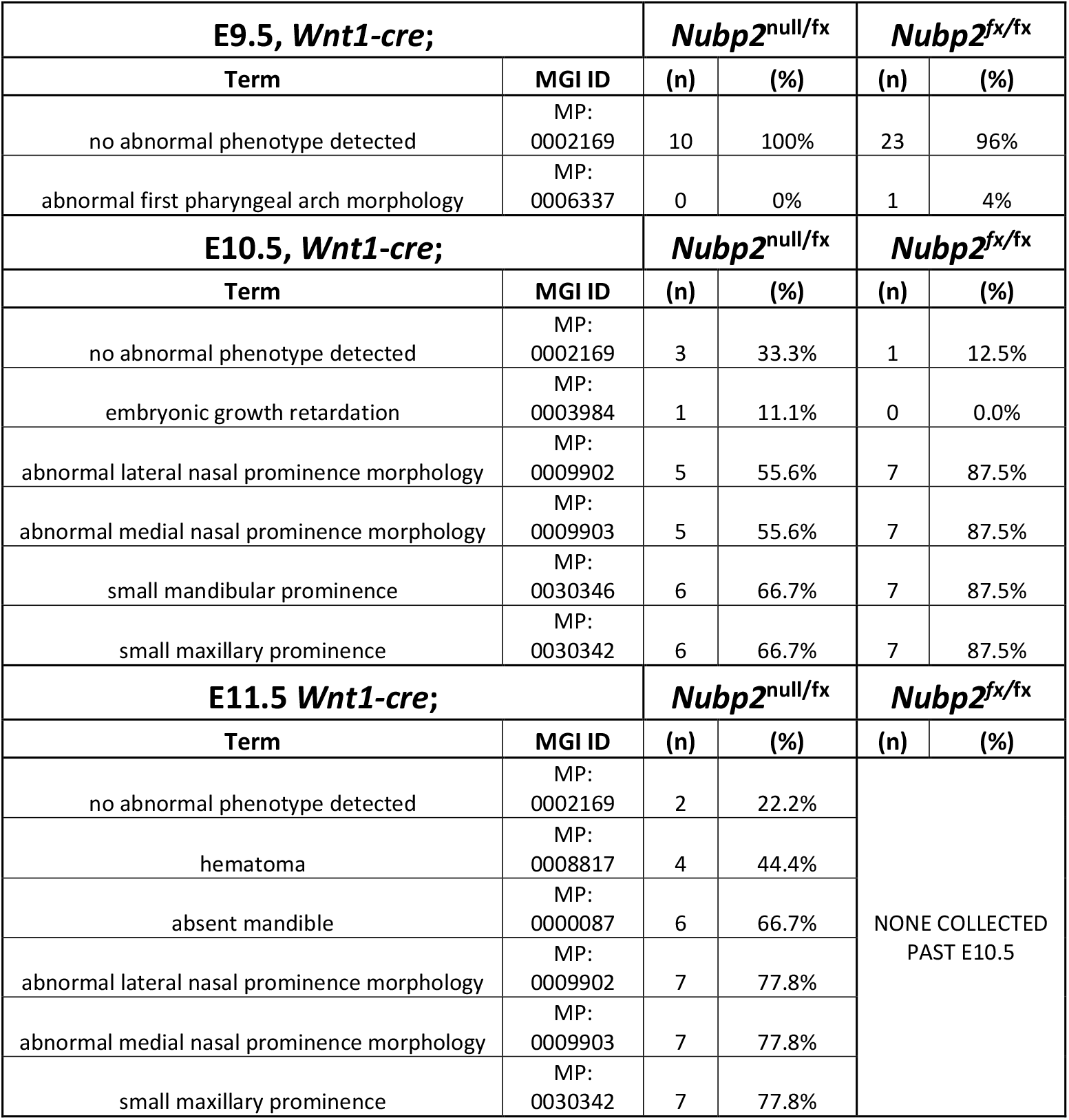
Comparison of phenotypes observed in *Wnt1-cre*; *Nubp2^null/fx^* and *Wnt1-cre*; *Nubp2^fx/fx^* embryos.

